# Human Lung Mast cell Diversity Across Anatomical Compartments Defined by Single-Cell Transcriptomics

**DOI:** 10.64898/2026.11.25.769139

**Authors:** Yitao Gong, Ionut Atanasoai, Daryl Boey, Anna-Karin Johnsson, Mamdoh Al-Ameri, Erik Sachs, Kasra Vali, Mikael Adner, Sven-Erik Dahlén, Jesper Säfholm, Joakim S. Dahlin, Gunnar P. Nilsson, Elin Rönnberg

**Author notes:** Shared senior authorship (Gunnar P. Nilsson, Elin Rönnberg). **Correspondence:** Elin Rönnberg Höckerlind, Address: L8:00 CMM, Visionsgatan 18, 171 64 Solna.

## Abstract

**Background:** Mast cells are key regulators of inflammatory lung diseases, including asthma. Yet, the heterogeneity of lung mast cells and their specialization in different lung compartments remain poorly defined. Traditional protease-based classifications fail to capture distinct mast cell states relevant to airway pathology, highlighting the need for high-resolution, compartment-resolved analyses.

**Objective:** To determine the heterogeneity of human lung mast cells in different lung compartments.

**Methods:** We performed single-cell RNA sequencing analysis of human lung mast cells from four anatomical compartments; bronchi, parenchyma, arteries and veins. Bioinformatical analysis was used to characterize mast cell heterogeneity and compartment-specific specialization.

**Results:** Five transcriptional distinct mast cell states (MC1-MC5) were identified. MC1 was characterized by stress response (e.g., *MT2A, MRPL14, SNN*), MC2 by broad immediate-early gene activation (e.g., *FOS, JUN*), and MC3 by a more selective activation with TGF-β response enrichment. MC4 displayed a pro-inflammatory effector phenotype enriched for IL-33-responses (e.g., *CCL2, CCL4, CCL4L2, IL13*), consistent with roles in airway inflammation. MC5 was associated with long non-coding RNAs (*MEG3, MEG8*). Mast cell states showed anatomical segregation with inflammatory cytokines enriched in bronchial and arterial mast cells while reduced in parenchymal mast cells. We further identified MC_TC_-like mast cells co-expressing *CTSG*, *CMA1* and *ADCYAP1*, characterized by enhanced metabolic programming.

**Conclusions:** This study provides a comprehensive single-cell transcriptomic atlas of human lung mast cells, revealing five distinct transcriptional states and compartment-specific functional specialization. These findings link mast cell heterogeneity to airway inflammation and provide a translational framework for targeting mast cells in lung diseases.

**Key Messages:**

- **Human lung mast cells comprise five transcriptionally distinct states (MC1–MC5)** with distinct activation, stress-response, and inflammatory programs, highlighting heterogeneity that is not captured by conventional protease-based classifications.
- **Mast cell states are distributed differently across anatomical compartments**, with inflammatory transcriptional programs enriched in bronchial and arterial mast cells but reduced in parenchymal mast cells, suggesting that the local tissue environment shapes mast cell specialization.

**Capsule summary:** Single-cell transcriptomics identified five transcriptionally distinct human lung mast cell states with compartment-specific specialization, revealing inflammatory enrichment in bronchial and arterial mast cells and providing a resource for future studies of inflammatory lung diseases.

## Introduction

Despite their established role in asthma and emerging relevance in other inflammatory lung diseases, the functional heterogeneity and compartment-specific organization of mast cells in the human lung remain largely unexplored. Mast cells are tissue-resident immune effector cells strategically positioned at host-environment interfaces, where they act as sentinels for immune surveillance and potent orchestrators of inflammatory responses (1). Mast cells are abundant in human lungs, where they comprise up to one third of the immune cells (2), and within the structurally complex lung, mast cells reside in highly divergent environments including the airways, parenchyma and pulmonary vasculature where they are exposed to unique microenvironmental cues that may shape and imprint their functional states. While their canonical role in driving asthma associated allergen induced bronchoconstriction via histamine and cysteinyl-leukotrienes is well documented, this traditional view captures only a limited aspect of mast cell biology. How mast cells and particularly how different mast cell phenotypes contribute to other features of the pathogenesis, e.g., airway remodeling, chronic inflammation and fibrosis, remain incompletely understood. This knowledge gap is particularly relevant given increasing evidence implicating mast cells in a broader spectrum of pulmonary pathologies, including, chronic obstructive pulmonary disease (COPD), and pulmonary fibrosis, where their anatomical localization and mediator profiles may influence disease progression and structural changes (3, 4). Consequently, resolving the distinct cellular diversity and compartment specific functional specialization of human lung mast cells (HLMC) is essential to fully understand their multifaceted roles in chronic lung disease.

A major conceptual limitation in the field is the reliance on a protease-based classification that divides human mast cells into tryptase-positive mast cells (MC_T_) and tryptase/chymase-positive mast cells (MC_TC_) expressing both tryptase, chymase, carboxypeptidase A3 and cathepsin G (5–7). While this framework has been instrumental in establishing foundational human mast cell biology, it is now over four decades old and insufficient to explain the diverse roles attributed to mast cells in lung diseases. Importantly, only a limited number of studies have interrogated HLMC heterogeneity beyond protease expression. Using immunohistochemistry, Andersson et al. described substantial heterogeneity within both MC_T_ and MC_TC_ populations across lung compartments, including differences in cell size and protein expression, highlighting that proteases content alone does not adequately capture HLMC diversity (8).

Recent advances in single-cell RNA sequencing (scRNA-seq) have enabled unprecedented resolution of cellular diversity, revealing states that are not identified by conventional protease-based classifications (9). In chronic rhinosinusitis with nasal polyposis (CRSwNP), scRNA-seq has revealed that mast cells exist along a transcriptional gradient rather than as discrete MC_T_ and MC_TC_ subsets, with tissue-specific microenvironments shaping mast cell phenotypes beyond shared protease expression (10). Nevertheless, comparable high-resolution analyses of mast cells in the human lung have been limited. Our previous scRNA-seq study of sorted HLMCs demonstrated extensively transcriptional variability beyond traditional protease-based classifications but lacked sufficient cell numbers to identify distinct subsets and mast cells from different anatomical lung compartments were not examined (11), suggesting that critical aspects of HLMC heterogeneity remain unresolved.

The lung is a structurally and functionally complex organ with multiple anatomical compartments including airways, parenchyma, and pulmonary vasculature, each providing unique microenvironmental cues that may imprint mast cells. Yet, a systematic characterization of mast cell heterogeneity across lung anatomical compartments is currently lacking. This represents a major knowledge gap, as mast cell functions are increasingly recognized to be context-dependent (12, 13), and compartment-specific mast cell states may differentially contribute to airway inflammation, vascular remodeling and fibrotic processes.

In this study, we address this gap by performing comprehensive single-cell transcriptomic profiling of HLMCs across four distinct anatomical compartments. Our analyses reveal five transcriptionally distinct mast cell populations and compartment-specific distribution patterns and functional signatures, providing new insights into the specialized roles of mast cells in human lung biology revealing functional programs that extend far beyond traditional protease-based classification.

## Materials and Methods

### Sample Collection and Processing

With approval from the Regional Ethical Review Board in Stockholm (ref. no. 2018/1819-31/1) and following informed consent, lung specimens were obtained from five patients undergoing lung surgery. Tissue samples were selected from macroscopically tumor-free regions located as distally as possible from the resected tumor and were immediately placed in ice-cold Krebs-Ringer physiological salt solution supplemented with 2.5 mM calcium chloride (Sigma-Aldrich) and 2.1 g/L sodium bicarbonate (Sigma-Aldrich). Within two hours of resection, the tissue was carefully dissected under microscopy into anatomically defined, compartment-enriched samples comprising intact bronchial rings, pulmonary arteries, pulmonary veins (only obtained from four donors), and lung parenchyma. Bronchial rings were prepared from small airways with an inner diameter of less than 2 mm and were identified by the presence of a clearly visible airway lumen and airway wall structure. Pulmonary arteries were identified as vessels with a relatively thick muscular wall, typically running in close association with bronchial branches, whereas pulmonary veins were distinguished by their thinner vessel wall and anatomical course separate from the bronchovascular bundle. To minimize cross-contamination between compartments, adherent connective tissue, blood clots, surrounding parenchyma, and adjacent vascular or airway structures were carefully removed during dissection. Parenchymal samples were collected from distal lung regions after careful removal of macroscopically visible airways and blood vessels.

### Single-Cell Isolation Protocol

Single-cell suspensions were prepared using an adapted protocol from Ravindran et al (14). Briefly, tissue from each anatomical compartment was cut into smaller pieces (2 mm) and placed in Epithelial Strip Buffer containing RPMI 1640 medium supplemented with 5% FCS, 5 mM EDTA, 15 mM HEPES, and 1 mM DTT for 30 minutes at 37°C. Thereafter, the tissue pieces (and cells) were washed with PBS to remove EDTA and DTT. Tissue pieces and cells were then placed in Enzyme buffer (RPMI 1640 with 15 mM HEPES, 0.2 mg/ml DNase I (Sigma-Aldrich) and 0.1 mg/ml Liberase TL (Sigma-Aldrich)) and incubated for 30 minutes at 37°C with magnetic stirring. The suspension was mechanically dissociated by passing through a syringe without needle 10 times, followed by passage through a 16G needle 10 times. Red blood cells were lysed using 5 ml ice-cold ACK buffer (Thermo Fisher) for 5 minutes. Mast cells were enriched using the CD117 MicroBead Kit (Miltenyi Biotec, #130-091-332) according to manufacturer’s instructions. Enriched cells were stained with anti-CD117-APC (Clone A3C6E2, Miltenyi Biotec, #130-091-733), anti-CD45-FITC (Clone HI30, BD Biosciences, #555482), and anti-CD14-APC-Cy7 (Clone M5E2, BioLegend, #301820). Cells from each anatomical compartment were stained with unique hashtag antibodies (BioLegend (#394631, #394633, #394635, #394637)) for 30 min. After washing, the cells were resuspended in PBS+ 2% FBS with EDTA containing DAPI (BD Biosciences, #564907). Mast cells were sorted on a BD FACSAria Fusion based on DAPI⁻ CD45⁺ CD14⁻ CD117^high^ expression profile. The hashtag-labeled cells from each compartment were combined and processed for 10x Genomics single-cell RNA sequencing with a target recovery of up to 20,000 cells.

### Single-Cell RNA Sequencing

Single-cell RNA sequencing libraries were prepared using the 10X Genomics Chromium Single Cell 3’ Reagent Kit v3 according to the manufacturer’s protocol. Libraries were sequenced on an Illumina NovaSeq platform and the bioinformatic analysis of the data was performed as detailed in the online repository.

## Results

### Identification of Five Mast Cell Transcriptional States in the Human Lung

To investigate mast cell heterogeneity across human lung compartments, mast cells isolated from four anatomical compartments, i.e., bronchi, arteries, veins and parenchyma, were individually hashtag-labelled, sorted, pooled, and sequenced together to minimize batch effects (Figure 1). Unsupervised clustering of single-cell RNA sequencing data from 19,232 mast cells identified five transcriptionally cell clusters (MC1-MC5) (Figure 2A), all of which were detected across samples from all five donors (see Figure E1 in the online repository). Each cluster exhibited a distinct transcriptional signature, characterized by its top differentially expressed genes, supporting heterogeneity among HLMCs (Figure 2B-C and Table E2 in the online repository).

**Figure 1.**
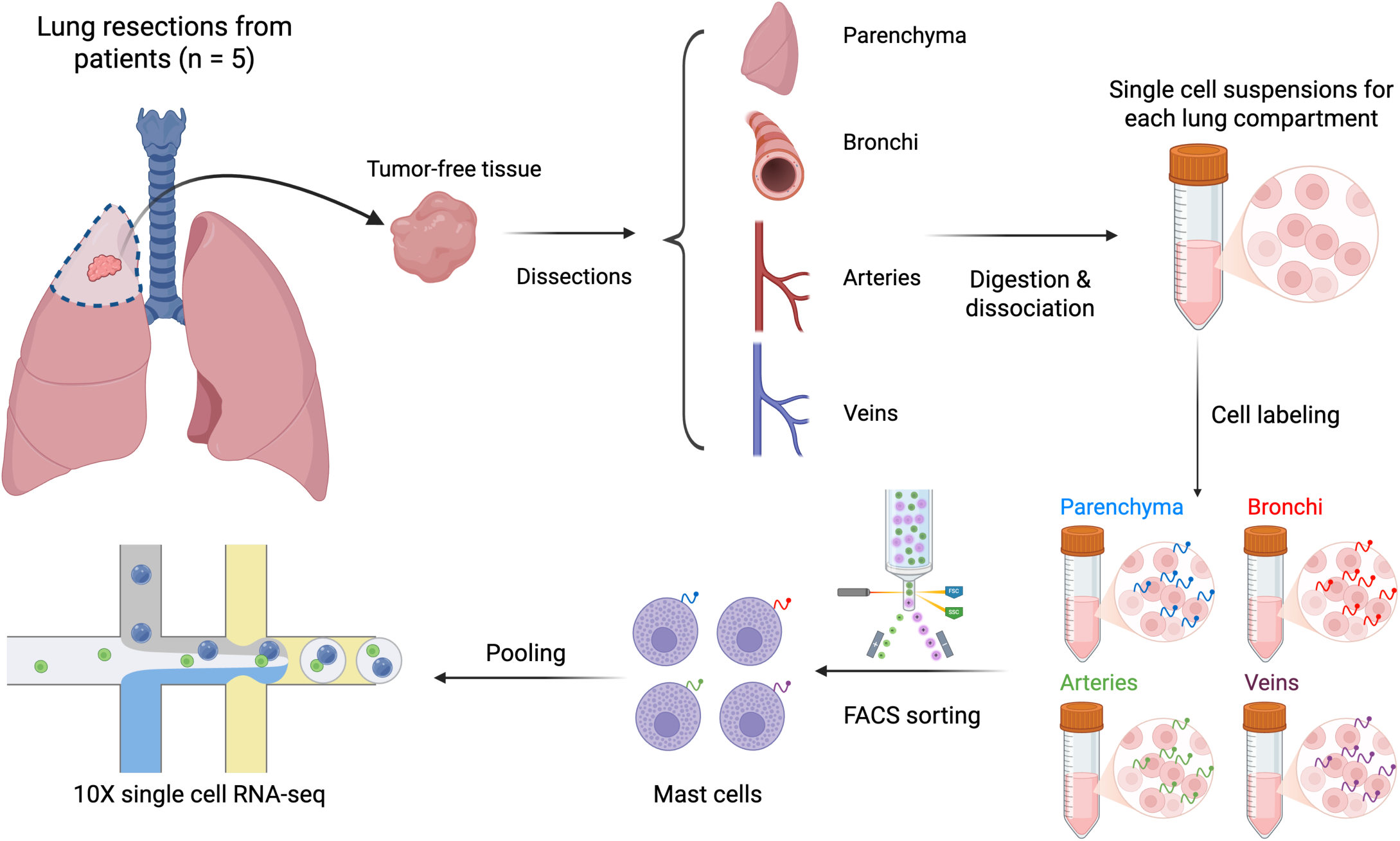
Single-cell RNA sequencing workflow for analyzing lung compartment-specific mast cell populations. Tumor-free tissue from 5 lung resections was dissected into four anatomical compartments: parenchyma, bronchi, arteries, and veins. Single-cell suspensions were generated by two-step digestion with epithelial stripping followed by DNase I and Liberase digestion. Cells from each compartment were labeled with compartment-specific hashtag antibodies. Mast cells were isolated by FACS, pooled, and scRNA-seq was performed using the 10x Genomics platform. Figure was created using BioRender.

**Figure 2.**
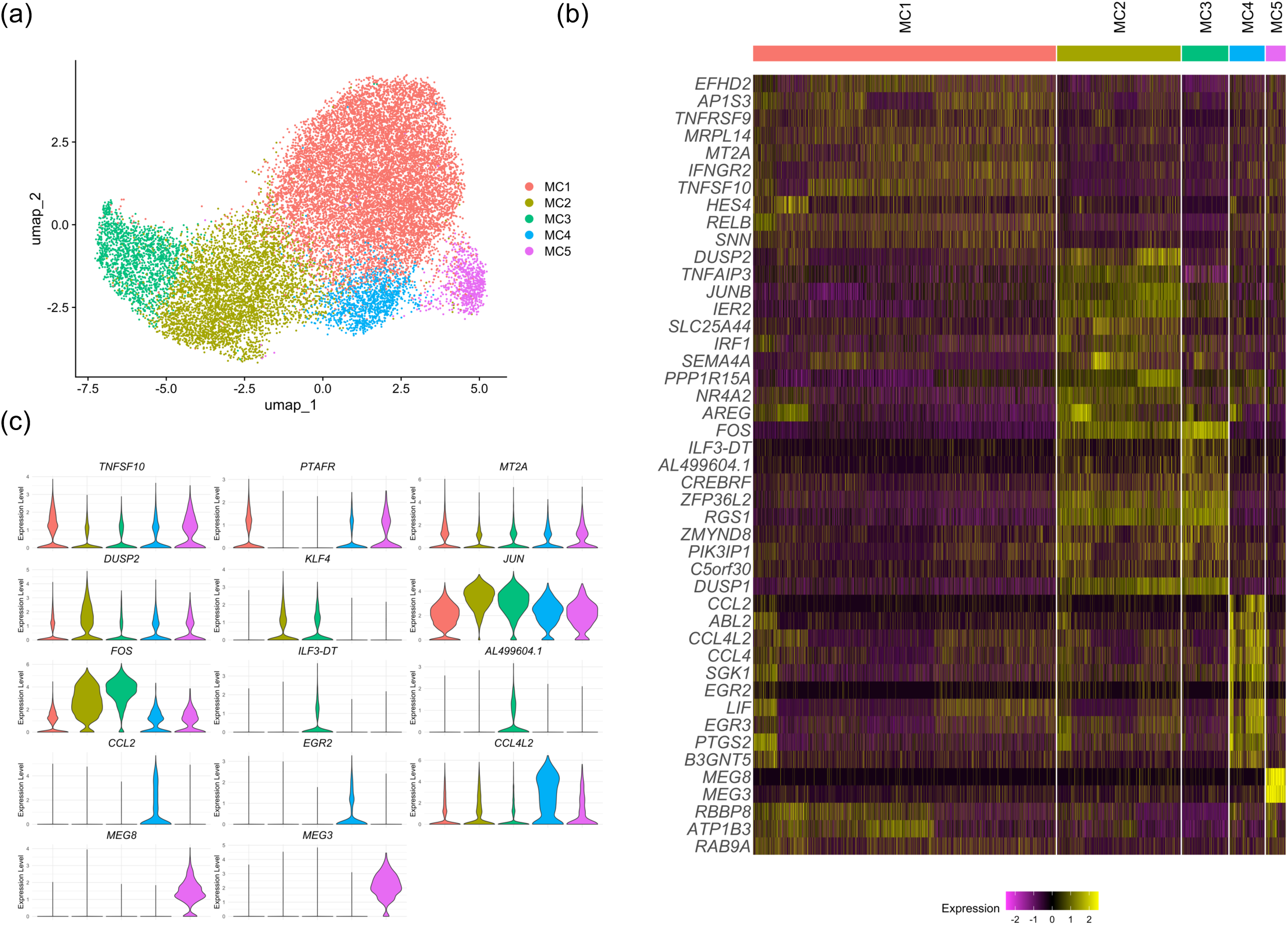
Identification and characterization of five distinct mast cell states. (A) UMAP visualization of single cell transcriptomics of 19,232 human lung mast cells showing five distinct clusters (MC1–MC5). (B) Heatmap of the 10 most enriched genes per cluster (five for MC5; see Methods and Table E2 of the online repository). (C) The three genes per cluster with the highest fold change relative to all other cells, among genes detected in more than 20% of the cells of that cluster. MC5 only had two such genes.

### Characterization of the Five Mast Cell Clusters

Characterization of the clusters revealed distinct transcriptional programs (Figure 2B-C and table E2 in the online repository). MC1 exhibited a stress-response-associated signature, characterized by increased expression of *MT2A, MRPL14* and *SNN*, together with the apoptosis-related gene *TNFSF10*. MC2 displayed a prominent immediate-early response signature, with elevated expression of transcription factors (*JUN*, *JUNB*), dual-specificity phosphatases (*DUSP2*), transcriptional regulators (*KLF4*, *NR4A2*), and the growth factor *AREG*, consistent with an activated transcriptional state. MC3 show more limited immediate-early response signature, characterized by *FOS* and *DUSP1*, together with the G-protein signaling regulator *RGS1* and the RNA-binding protein *ZFP36L2*. In contrast, MC4 displayed a strong pro-inflammatory transcriptional profile, with enrichment of cytokine signaling pathways (see Figure E2-E3 in the online repository) due to high expression of cytokines and chemokines. Finally, MC5 closely resembled MC1 but exhibited selective expression of *MEG3*and *MEG8*, suggesting potential differences in non-coding RNA-mediated gene regulation.

Analysis of functional gene expression patterns across the five MC states revealed distinct expression profiles among key effector molecules in three functional categories (Figure 3). MC4 exhibited high expression of multiple chemokines (*CCL4L2, CCL2, CCL4*) and interleukins (*IL13, IL5*), while MC3 showed higher expression of *CCL28, CCL16* and *IL18*. Prostaglandin related genes, *PTGS2* and *PLA2G4A,* were highly expressed in MC4, whereas MC3 preferentially expressed other enzymes involved in eicosanoid production including *ALOX5, LTC4S*, *HPGDS* and *PTGS1*. Transcription factor expression also differed between the clusters, with MC2 showing particularly high expression of immediate-early response genes (*FOS, FOSB, JUN, JUNB*), whereas MC3 was characterized by increased expression of interferon-associated transcription factors, including *IRF2, IRF3, IRF9*, and *STAT1*.

**Figure 3.**
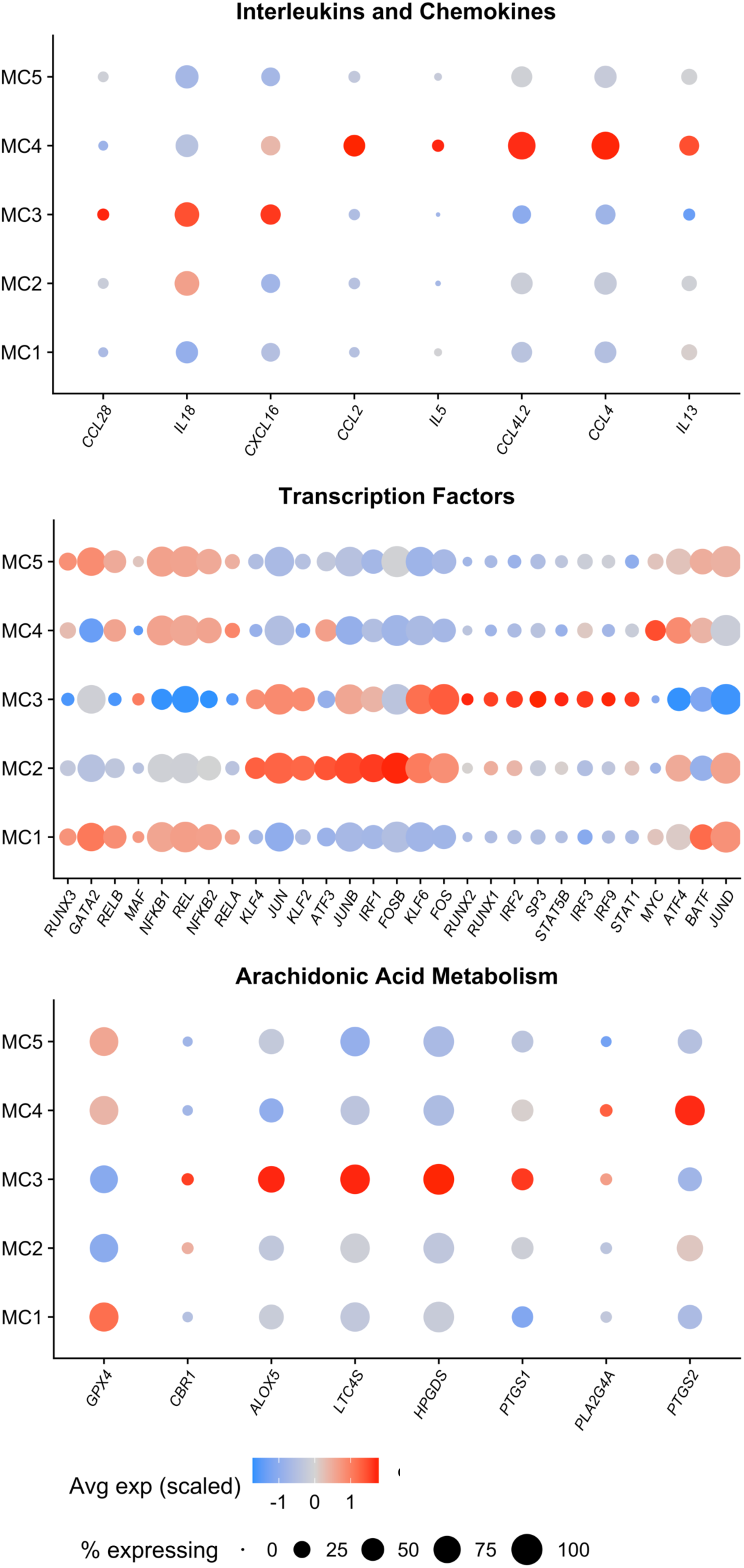
Functional genes across the five mast cell states. Dot plots of functional genes across three categories: Interleukins and Chemokines, Transcription Factors, and Arachidonic Acid Metabolism. Genes were selected from all differentially expressed genes among the five clusters based on HUGO Gene Nomenclature Committee annotations and restricted to genes detected in more than 10% of the cells of the cluster in which they were identified. Dot size indicates percentage of expressing cells; color represents average expression.

### Expression of Canonical Mast Cell Markers

Despite the transcriptional differences among the five clusters, all MC clusters retained expression of nine canonical mast cell genes, although expression levels varied between clusters for certain genes (Figure 4A). The mast cell protease genes *TPSAB1*, *TPSAB2*, and *CPA3* were broadly expressed across MC clusters, with *TPSAB1* showing considerable intra-cluster variation. MC3 exhibited higher expression of *KIT* (CD117, the stem cell factor receptor) and *MS4A2* (encoding the β-chain of the high-affinity IgE receptor, FcεRI), as well as *HPGDS*, an essential enzyme involved in prostaglandin D₂ synthesis. In contrast, *HDC*, encoding histidine decarboxylase, the rate-limiting enzyme in histamine biosynthesis, was more highly expressed in MC1. Together, these findings demonstrate that the five transcriptional clusters maintain a common mast cell identity while exhibiting cluster-specific differences in the expression of genes associated with mediator synthesis and receptor signaling.

**Figure 4.**
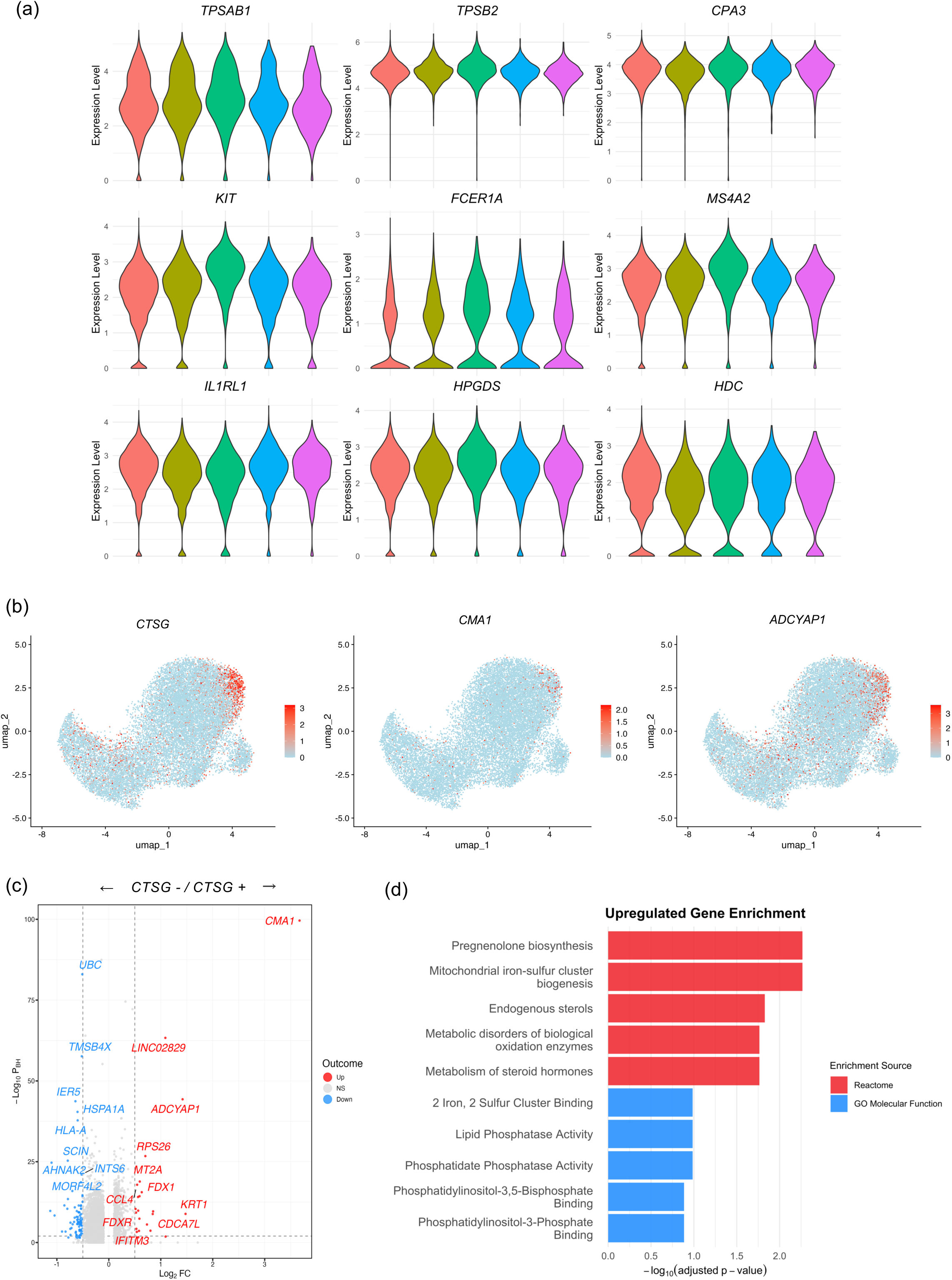
Expression of canonical mast cell markers and characterization of MC_TC_-like cells. (A) Expression of nine canonical mast cell markers across clusters. (B) UMAP feature plots showing co-expression of *CTSG, CMA1*, and *ADCYAP1* defining MC_TC_-like cells. (C) Volcano plot of differentially expressed genes between CTSG-positive and CTSG-negative cells. (D) Gene set enrichment analysis showing metabolic pathway enrichment in CTSG-positive cells.

### Identification and Characterization of MC_TC_-like Cells

To establish the connection between our single-cell transcriptomically defined clusters and traditional histochemically defined mast cell subtypes, we examined expression of proteases that define the MC_TC_ phenotype: chymase (*CMA1*), carboxypeptidase A3 (*CPA3*), and cathepsin G (*CTSG*). While *CPA3* was broadly expressed (Figure 4A), *CTSG* and *CMA1* were enriched in a distinct region of the UMAP projection, where their expression overlapped with that of *ADCYAP1* (Figure 4B).

Differential gene expression analysis comparing *CTSG*-positive and *CTSG*-negative populations revealed distinct molecular signatures (Figure 4C). *CTSG*-positive cells exhibited higher expression of for chymase (*CMA1*), structural proteins (*KRT1*), neuropeptides (*ADCYAP1*), and metabolic regulators, while *CTSG*-negative cells were characterized by elevated expression of stress response genes (*HSPA1A*), cytoskeletal components (*TMSB4X*) and antigen presentation molecules (*HLA-A*). Gene set enrichment analysis further highlighted the metabolic specialization of *CTSG*-positive cells, revealing significant enrichment in pathways related to steroid hormone biosynthesis pathways, including pregnenolone biosynthesis, mitochondrial processes, and steroid hormone metabolism (Figure 4D).

### Cytokine-Responsive Signatures Define Functional Polarization

To assess cytokine-associated transcriptional states among the mast cell clusters, we applied published gene signatures derived from mast cells exposed to IFN-γ, IL-33, IL-4/IL-13, and TGF-β (15). The IL-33-responsive signature was most pronounced in MC4, suggesting enhanced transcriptional responsiveness to this epithelial-derived alarmin in this cluster. IFN-γ- and TGF-β-responsive signatures were enriched in both MC2 and MC3, with the TGF-β signature showing particularly strong enrichment in MC3. In contrast, IL-4/IL-13-responsive signatures showed limited variation between clusters. This may reflect a shared capacity to respond to type 2 cytokines or limited differences in their exposure to these signals in the tissue (Figure 5).

**Figure 5.**
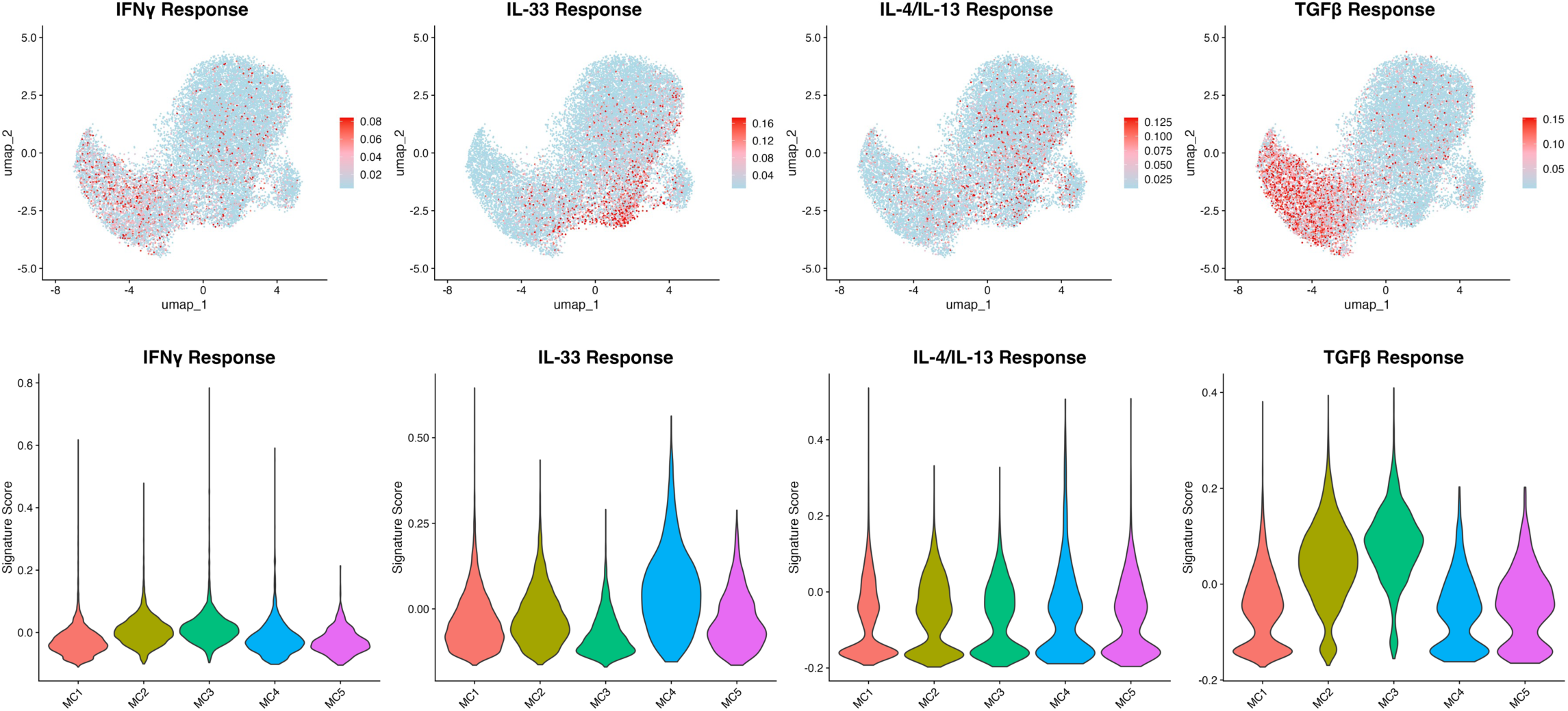
Cytokine-responsive signatures define functional polarization. Enrichment of cytokine response genes from Tontini et al. (15) in UMAP projections and violin plots quantifying score distributions across the five mast cell clusters (MC1-MC5).

### Cross-Dataset Validation and Reproducibility

To validate our findings within the broader context of mast cell heterogeneity, we performed comprehensive cross-dataset comparisons with previously published single-cell datasets. Visualization of the MC1–MC6 mast cell signature scores defined by Tauber et al. (13), who analyzed mast cells from multiple anatomical sites in the body, on our UMAP revealed that their MC1–MC5 signatures were associated with distinct regions, whereas the MC6 signature showed no clear regional enrichment (Figure 6A). Notably, their MC1 signature aligned with our MC3 cluster, while their MC3, characterized by high expression of *CTSG* and *ADCYAP1* expression, overlapped with our *CTSG-*positive region.

**Figure 6.**
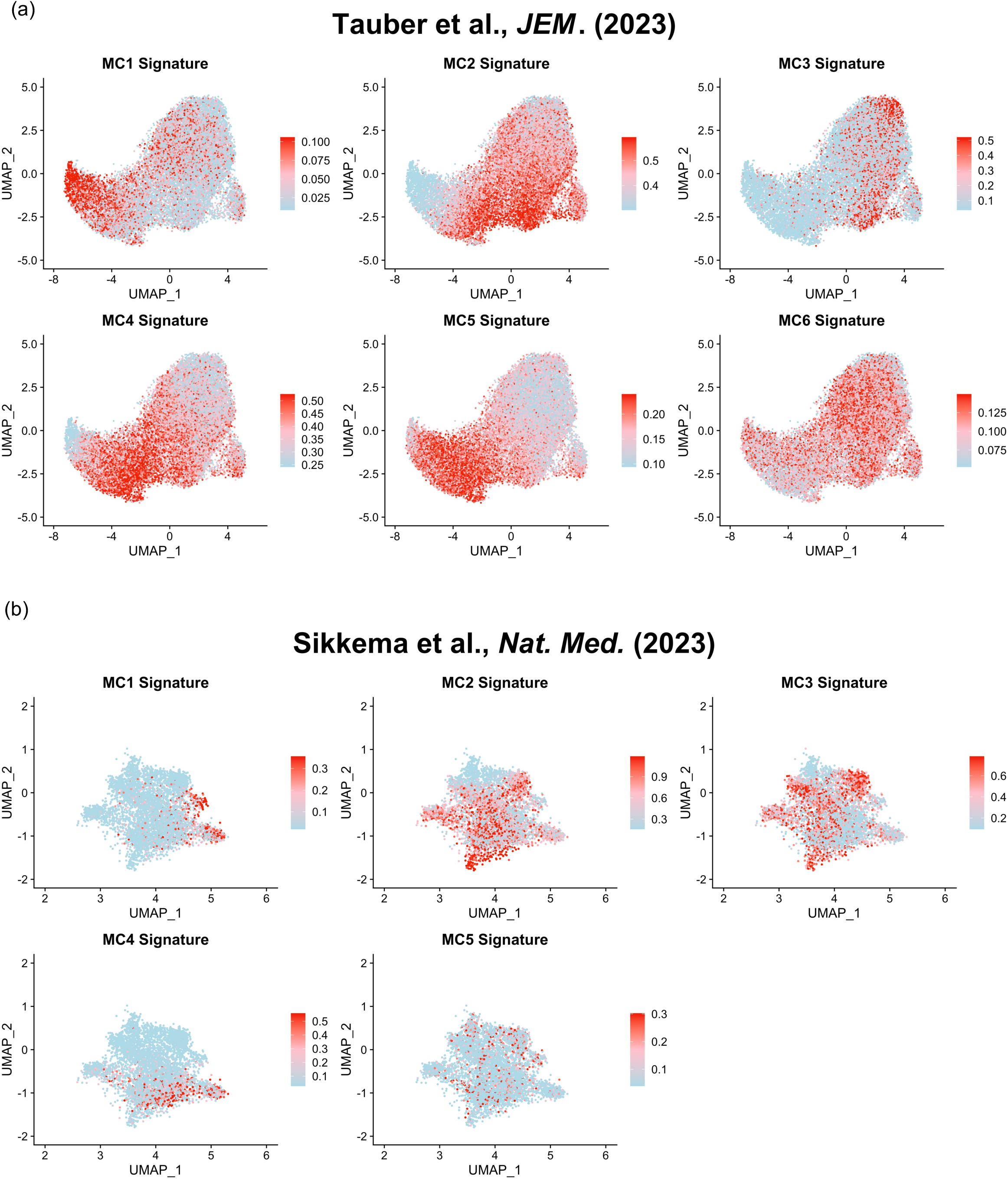
Cross-dataset validation of mast cell subtypes. (A) Visualization of the MC1–MC6 mast cell signature scores from the Tauber et al. study on our dataset. (B) Visualization of our MC1-MC5 signatures on Sikkema et al. lung atlas dataset. Signatures comprised the 20 most significant marker genes per cluster (10 for MC5; see Table E2 of the online repository). Expression intensity shown by color scale (blue = low, red = high), scaled between the 10th and 90th percentile within each panel.

Conversely, visualization of our MC1–MC5 signature scores (see table E2 and Figure E4 in the online repository) on re-analyzed mast cells from the human lung cell atlas (16) showed that our MC1-MC4 were associated with distinct regions of the atlas UMAP (Figure 6B). However, the MC5 cluster, characterized by expression of the long non-coding RNAs, *MEG3* and *MEG8*, lacked a clear counterpart in external datasets. This suggests either a potentially lung-specific transcriptional program or underrepresentation of this population in previous datasets, which did not employ mast cell enrichment prior to single-cell analysis.

### Compartment-Specific Distribution and Functional Specialization

Analysis of HLMCs from the four anatomical lung compartments, i.e., bronchi, arteries, veins and parenchyma, revealed distinct compartment-specific distributions of the five mast cell clusters (Figure 7A). MC4 was enriched in mast cells localized in arteries and bronchi and underrepresented in the parenchyma. MC2 and MC3 were also enrichment in the bronchial compartment, whereas MC1 and MC5 were more evenly distributed across compartments.

**Figure 7.**
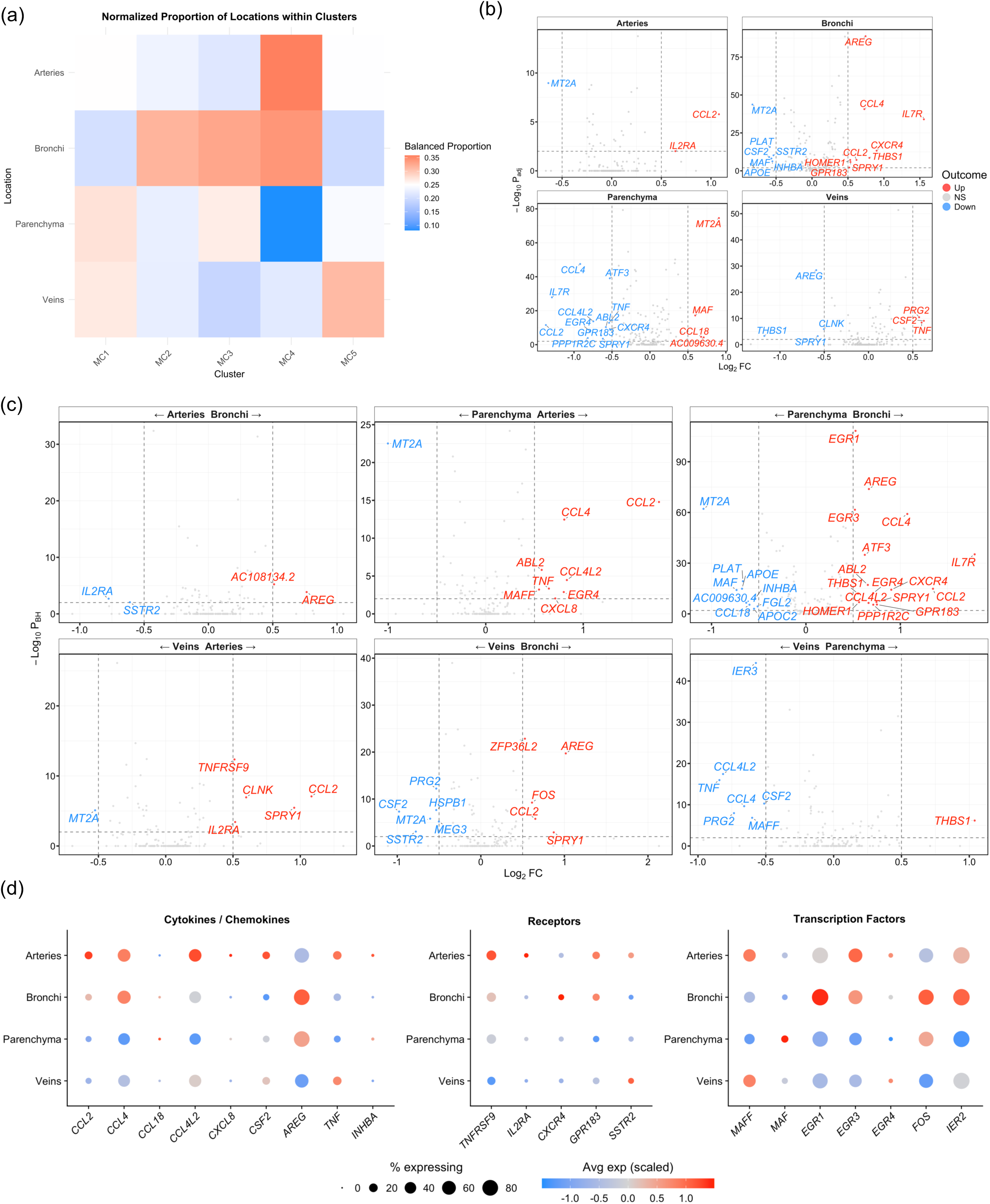
Compartment-specific distribution and functional specialization of mast cells. (A) Heatmap showing normalized proportions of MC1-MC5 across anatomical compartments. (B) Volcano plots of differentially expressed genes between mast cells from each anatomical compartment compared to all other cells. (C) Pairwise comparisons between compartments showing location-specific transcriptional signatures. Genes with |log₂FC| > 0.5 and adjusted p < 0.01 are highlighted. (D) Dot plots of differentially expressed genes across compartments in three categories.

Differential gene expression analysis identified compartment-specific transcriptional programs associated with each anatomical location (Figure 7B-D). Bronchial mast cell populations showed selective enrichment for immediate early transcription factors (*FOS, EGR1, EGR3* and *IER2)* together with an inflammatory signature characterized by elevated expression of multiple mediators (*AREG, CCL4, CCL2*) and chemokine receptor *CXCR4*. Arterial and venous mast cells shared several transcriptional features, including elevated expression of *TNF, CSF2, SST2* and the transcription factor *MAFF*. However, arterial mast cells demonstrated a more pronounced inflammatory signature, with elevated expression of multiple chemokines (*CCL2, CCL4L2, CCL4, CXCL8*), as well as cytokine receptors (*TNFRSF9, IL2RA*) and G-protein coupled receptor *GPR183*. In contrast, parenchymal mast cells generally exhibited lower expression of inflammatory mediators, including *CCL4, CCL2, CCL4L2,* and *TNF*.

## Discussion

This study provides the first comprehensive single-cell transcriptomic analysis of human lung mast cells across major anatomical compartments and demonstrates that HLMC diversity is substantially greater than accounted for by traditional protease-based classifications. By analyzing 19,232 purified HLMCs, we identified five transcriptional clusters (MC1-MC5). These findings provide a revised conceptual framework for mast cell biology in the human lung by building upon recent insights showing that classical definitions vastly underestimate their true functional diversity (9, 17).

The five MC clusters identified exhibited distinct transcriptional and functional programs. MC4 cells displayed a strong pro-inflammatory signature characterized by IL-33 response enrichment and elevated chemokine and cytokine expression (Figure 3 and 5), preferentially localized to bronchial and arterial compartments (Figure 7A) where rapid immune responses are essential. In contrast, MC1 cells expressed stress-response genes (Figure 2), and had lower expression of cytokines/chemokines (Figure 3), suggesting a specialized role in stress response. MC2 and MC3 exhibited distinct transcriptional activation profiles, with MC3 notably enriched for TGF-β–response enrichment (Figure 5) and arachidonic acid metabolism genes (Figure 3). Finally, MC5 was defined by high expression of long non-coding RNAs (lncRNAs) MEG3 and MEG8 (Figure 2), suggesting epigenetic regulation (18). Cross-dataset validation demonstrates the robustness and reproducibility of the identified mast cell states MC1-MC4 across two independent datasets (13, 16), confirming that these mast cell states represent fundamental aspects of human mast cell biology rather than technical artifacts (Figure 6). The lncRNA-enriched MC5 population lacked clear correspondence in external datasets suggesting either a rare or context-dependent state, not consistently captured across studies. This comprehensive validation is particularly important given the rapid expansion of mast cell scRNA-seq studies and establishes a foundation for reproducible findings across different experimental contexts. ScRNAseq of mast cells from nasal polyps in CRSwNP, a T2 cytokine-mediated inflammatory disease, revealed the existence of a proliferative mast cell cluster (10), while in our dataset proliferative mast cells are distinctly lacking. We can barely detect any mRNA expression of the proliferation marker Ki-67 (*MKI67*, data not shown), nor have we been able to detect the protein by flow cytometry (19), indicating that to the contrary to mast cell in the nasal polyps, HLMCs are primarily long-lived non-proliferative cells.

The identification of compartment-specific mast cell signatures supports the concept that these cells undergo context-specific modifications of their effector properties in response to local microenvironmental signals (12). Bronchial mast cells, enriched for inflammatory mediators such as *CCL4* and *AREG* (Figure 7B-D), appear specialized for rapid immune responses against inhaled pathogens and allergens, with *AREG* expression in particular contributing to epithelial remodeling and goblet cell hyperplasia in asthma (20). In contrast, parenchymal populations show lower expression of inflammatory mediators (Figure 7D), suggesting that these cells are less prone to induce inflammatory responses. The distinct profiles of arterial and venous mast cells, with arterial populations showing a broader inflammatory capacity (Figure 7C-D), further demonstrate the sophisticated adaptation of mast cells to their specific anatomical niches. A recent study using spatial transcriptomic analysis (21) has provided validation for compartment-specific adaptations, demonstrating that bronchial mast cells expressing *AREG* and *CCL4* are strategically positioned within inflammatory hubs at epithelial-subepithelial interfaces. Within these hubs, mast cells are in close proximity to IL-33-producing basal cells and chemokine-secreting endothelial cells, creating localized signaling networks that can sustain the inflammatory phenotype we observed in bronchial mast cells. This spatial organization explains how local microenvironmental signals drive the context-specific modifications we identified through transcriptional profiling.

Classically, mast cells are subdivided into MC_T_ and MC_TC_ based on their expression of mast cell proteases, with MC_T_-cells expressing tryptase and MC_TC_ in addition expressing chymase (*CMA1*), CPA3 and cathepsin G (*CTSG*). We also identified a *CTSG*⁺ *CMA1*⁺ MC_TC_-like population with unique metabolic programming including steroid hormone biosynthesis pathways (Figure 4B-D). TGF-β has been identified as a key driver of the MC_T_ transcriptome in nasal polyps as well as *in vitro* peripheral blood derived mast cells (22). Concordantly, we see an enrichment of TGF-β response genes in the opposite side of the UMAP (Figure 5) to our MC_TC_-like population (Figure 4B), suggesting that TGF-β may also play a role in the maintenance of the MC_T_ cells in human lungs. The *ADCYAP1* expression in the MC_TC_-like population is particularly noteworthy, as *ADCYAP1* encodes pituitary adenylyl cyclase-activating polypeptide (PACAP), a neuropeptide involved in airway smooth muscle relaxation and bronchomotor regulation (23). The enrichment of *ADCYAP1* in *CTSG+* mast cells positions these mast cells for local PACAP release, potentially contributing to airway smooth muscle modulation. PACAP can also activate mast cells through the MRGPRX2 receptor (24), this receptor is however not expressed by HLMCs, nor do they respond to MRGPRX2 agonists (25, 26), making it unlikely that PACAP could have an autocrine function on mast cells in the lung. Our CTSG^+^ population aligns with Tauber et al.’s identification of a population characterized by high *CTSG* and *ADCYAP1* expression (13), reinforcing the biological significance of these subtypes and our data provides new insights into the functional specialization of this traditionally defined subset.

Several limitations of the study merit consideration. First, although tumor-free tissue was utilized, lung cancer–associated effects cannot be excluded. Second, dissociation-induced transcriptional changes potentially contributed to the high immediate early gene signatures observed in MC2 and bronchial mast cells. However, all tissues were treated the same and cells were hashtag-labeled and processed together to minimize batch effects, thus, these changes potentially reflect differential susceptibility to the processing stress rather than biological activation alone (27). Third, while transcriptomic specialization was clear, functional validation remains essential to determine physiological and pathological relevance.

Future studies should apply spatial transcriptomics to precisely map mast cell states within lung architecture, perform functional perturbation experiments to test predicted pathways, and define how these mast cell states change during disease progression. The enrichment of inflammatory MC4 cells in bronchial and arterial compartments points to potential involvement in airway hyperresponsiveness and vascular inflammation.

In summary, this work establishes an anatomical compartment-resolved framework of HLMC heterogeneity, identifying five transcriptionally distinct mast cell states with discrete functional attributes. These findings move the field beyond classical protease-based mast cell taxonomy, highlighting the combined influence of intrinsic transcriptional programming and anatomical location on mast cell function. This atlas provides a foundation for developing state- and compartment-specific therapeutic strategies targeting mast cells in respiratory disease.

## Author Contributions

E.R., J.S.D. and G.N. conceived and designed the study. M.A., E.S., K.V. and J.S. provided the human lung tissue and J.S. performed the tissue dissection of anatomical compartments. Y.G. and E.R. performed the experiments. Y.G., D.B., and I.A. conducted bioinformatics analyses. A-K. J., M.A., S-E. D., J.S. and J.S.D. provided technical expertise and contributed to data interpretation. G.N. and E.R. supervised the project. Y.G., G.N., and E.R. wrote the manuscript with input from all authors.

## Acknowledgments

We thank the patients who provided tissue samples for this study.

## Data Availability

All raw sequencing data and processed gene expression matrices are deposited in the Gene Expression Omnibus (GEO) database, accession number GSE326140 (will be made public upon publication). Analysis code and protocols are available upon reasonable request.

## Declaration of generative AI and AI-assisted technologies in the manuscript preparation process

During the preparation of this work the authors used Claude Sonnet 4.5 (Anthropic) and ChatGPT to polish the language. After using the tools, the authors reviewed and edited the content as needed and takes full responsibility for the content of the publication.

## Abbreviations

(COPD): chronic obstructive pulmonary disease
(HLMC): human lung mast cells
(MC_T_): tryptase-positive mast cells
(MC_TC_): tryptase/chymase-positive mast cells
(scRNA-seq): single-cell RNA sequencing
(CRSwNP): chronic rhinosinusitis with nasal polyposis
(PACAP): pituitary adenylyl cyclase-activating polypeptide

