## Supplementary material for "Human Lung Mast cell Diversity Across Anatomical Compartments Defined by Single-Cell Transcriptomics": Supplemetal File

*Shared senior authorship.

**Correspondence**

Elin Rönnberg Höckerlind

Address: L8:00 CMM, Visionsgatan 18, 171 64 Solna

Phone number: +46 768117819

#### Materials and methods

*Bioinformatics Analysis*

*Software and versions*: Unless stated otherwise, analyses were performed in R 4.5.3 with Bioconductor 3.22, using Seurat 5.4.0. The initial per-donor processing and donor integration were performed with Seurat 5.0.2, as recorded in the saved objects. Additional packages were multimode 1.5, MAST 1.36.0, leidenbase 0.1.37, fgsea 1.36.2, ReactomePA 1.54.0 and enrichR 3.4. Unless another package is named, all functions referred to below are Seurat functions.

*Initial Processing*: Raw sequencing data were processed using Cell Ranger software (10X Genomics): Cell Ranger v5.0.1 (donor 1), v7.0.1 (donors 2-3), and v7.1.0 (donors 4-5). Reads were aligned to the GRCh38-2020-A human reference transcriptome without intron inclusion. Cell barcodes and unique molecular identifiers (UMIs) were demultiplexed, and gene expression matrices were generated for downstream analysis.

*Hashtag Oligo Demultiplexing:* Anatomical compartment assignment was performed using a custom manual demultiplexing algorithm implemented in R. The algorithm assumed bimodal distribution of hashtag oligo (HTO) counts and identified threshold values by determining the dip between two modes using the locmodes() function from the multimode R package (v1.5). Upper and lower thresholds were calculated using quartile-based outlier detection methods (75th percentile + 1.5×IQR for lower bound, 25th percentile - 1.5×IQR for upper bound). Only singlet cells (assigned to a single compartment) were retained for further analysis.

*Quality Control and Filtering:* Quality control metrics including mitochondrial gene content, total UMI counts, and number of detected features were calculated for each cell using Seurat. Donor-specific filtering criteria were applied based on quantile analysis to remove low-quality cells and potential doublets (see Table E1 in the online repository).  Cells exceeding the donor-specific mitochondrial cutoff (range ≤8% to ≤18%) or falling outside the donor-specific lower and upper bounds for UMI counts or detected genes were excluded to remove low-quality cells and potential doublets.  Genes detected in fewer than 3 cells were removed from the analysis.

*Data Integration and Normalization:* Individual donor datasets were processed separately with the Seurat functions NormalizeData (normalization.method = "LogNormalize", scale.factor = 10,000) and FindVariableFeatures (selection.method = "vst", nfeatures = 500 per donor). Integration anchors were identified with FindIntegrationAnchors using the variable features shared by all five donors, and the donors were integrated with IntegrateData. The integrated assay was scaled with ScaleData.

*Dimensionality Reduction and Clustering:* Principal component analysis was performed with RunPCA (npcs = 15), and the first 15 principal components were used for both RunUMAP and FindNeighbors. Clusters were defined with the Seurat function FindClusters, using the Leiden algorithm (argument algorithm = 4, implemented through the leidenbase package) at resolution = 0.4 with random.seed = 1.

*Cell Type Validation:* Mast cell identity was confirmed by expression of canonical markers including *TPSAB1*, *CPA3*, *KIT*, and *FCER1A*. Non-mast cell populations were identified by expression of lineage-specific markers and excluded from downstream analysis.

*Differential Expression Analysis:* Differentially expressed genes between clusters and anatomical compartments were identified with the Seurat functions FindAllMarkers and FindMarkers, using the MAST hurdle model (argument test.use = "MAST") and including donor as a latent variable (argument latent.vars = "Donor").

*Marker Gene Selection for heatmap:* The top 50 genes per cluster are listed in Table *E*2. For the Figure 2B heatmap, each gene was assigned to the cluster where its scaled expression was highest, and the 10 genes per cluster with the largest difference from the next highest cluster were shown, restricted to genes detected in more than 25% of cells in that cluster. Genes of the MTRNR2L family, which are nuclear pseudogenes of mitochondrial rRNA and track mitochondrial content rather than cell identity, were excluded, both here and from the cluster signatures. No significance threshold was applied, so the panel shows relative enrichment rather than cluster-exclusive expression. When each displayed gene was compared between its cluster and the next highest cluster (Wilcoxon rank sum test with Bonferroni correction), all genes shown for MC2, MC3 and MC4, and MEG3 and MEG8 for MC5, were significant, whereas the remaining MC5 genes and the MC1 genes were not.

*Functional Enrichment Analysis:* Over-representation analysis of cluster marker genes was performed with ReactomePA (v1.54.0) and enrichR (v3.4) (GO Molecular Function and Biological Process; significant at adjusted $p < 0.05$). Per-cluster gene set enrichment analysis (GSEA) was additionally performed with fgsea (v1.36.2) against the MSigDB Hallmark and Reactome collections, ranking genes by signed significance ($\text{sign}(\log_2 FC) \times -\log_{10} p$) from the same MAST model (donor as a latent variable); up-regulated pathways (NES > 0, adjusted $p < 0.05$).

*Signature Score Analysis:* Module scores for signature enrichment were calculated with the AddModuleScore function. Cytokine response signatures were extracted from the differentially expressed genes reported by Tontini et al. (E1). Specifically, for each cytokine condition (IFNγ, IL-33, IL-4+IL-13, TGFβ) we used their reported DEGs, defined at FDR < 0.05 and $\log_2$ fold change > 5 (51, 39, 14, and 15 genes, respectively). These gene sets were scored with AddModuleScore and aggregated per mast-cell cluster.

For cross-dataset validation we projected signatures in both directions. (i) Forward: marker genes of the six mast-cell clusters (MC1–MC6) reported by Tauber et al. (E2) were scored on our cells with AddModuleScore. (ii) Reciprocal: the top 20 marker genes of each of our five clusters (MC1–MC5) were scored on external mast cells. For the Human Lung Cell Atlas published by Sikkema et al. (E3), we took the published mast-cell subset and its integrated scANVI embedding, computed a new UMAP from that embedding, and overlaid the MC1–MC5 module scores on it; the resulting layout therefore differs from the UMAP shown on the atlas portal. Several signature genes, mostly unnamed lncRNAs, are not annotated in the atlas and were therefore absent from the corresponding scores (8 of 20 genes for MC3, 1 of 20 for MC4 and 2 of 10 for MC5). As a positive control, the same signatures scored on our own UMAP were each enriched in their corresponding cluster (see Figure E4 in the online repository).

*Anatomical Distribution Analysis:* To compare the distribution of mast-cell clusters across anatomical compartments while accounting for differences in cell recovery, we first calculated, within each compartment, the proportion of cells assigned to each cluster. These proportions were then normalized within each cluster to sum to one across compartments, so that the resulting values reflect the relative enrichment of each cluster across compartments independently of the number of cells recovered. Balanced proportions were computed in base R (table, ave) and visualized as heatmaps with ggplot2.

Statistical Analysis: Analyses were performed in R (version 4.5.3). Differential expression between clusters and between anatomical compartments was assessed with the MAST hurdle model (Seurat FindAllMarkers/FindMarkers, test.use = "MAST"), including donor as a latent variable to account for inter-donor variation; genes with a Benjamini–Hochberg-adjusted p-value $< 0.05$ were considered significant. Summary quantities (e.g., module scores and cluster proportions) were compared using the Kruskal–Wallis test (multiple groups) and the Mann–Whitney U test (pairwise comparisons).

### Online Repository Figures

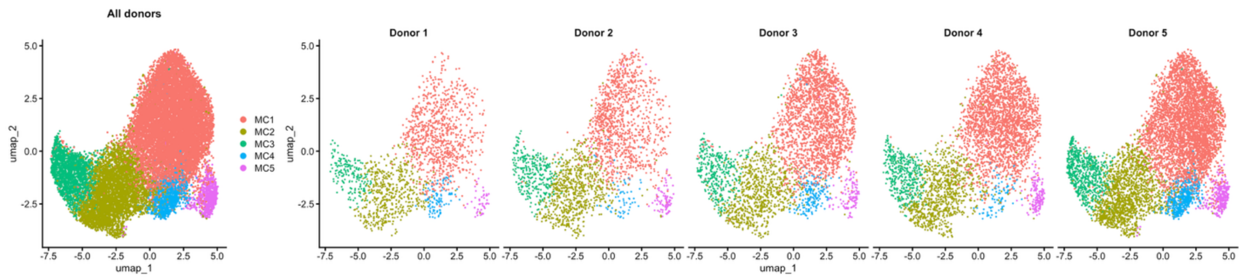


**Figure E1.** Contribution of the different donors to the UMAP**.**

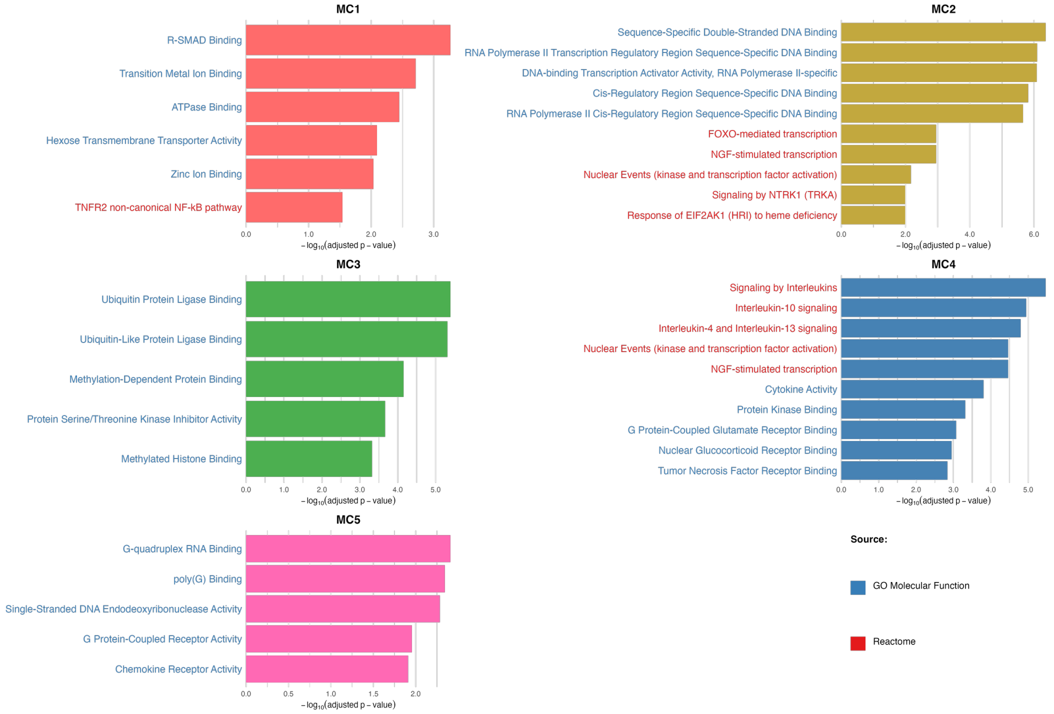


**Figure E2.** Over-representation analysis of cluster marker genes (GO Molecular Function and Reactome). Bars show −log10 adjusted p-value; colour indicates the source database.

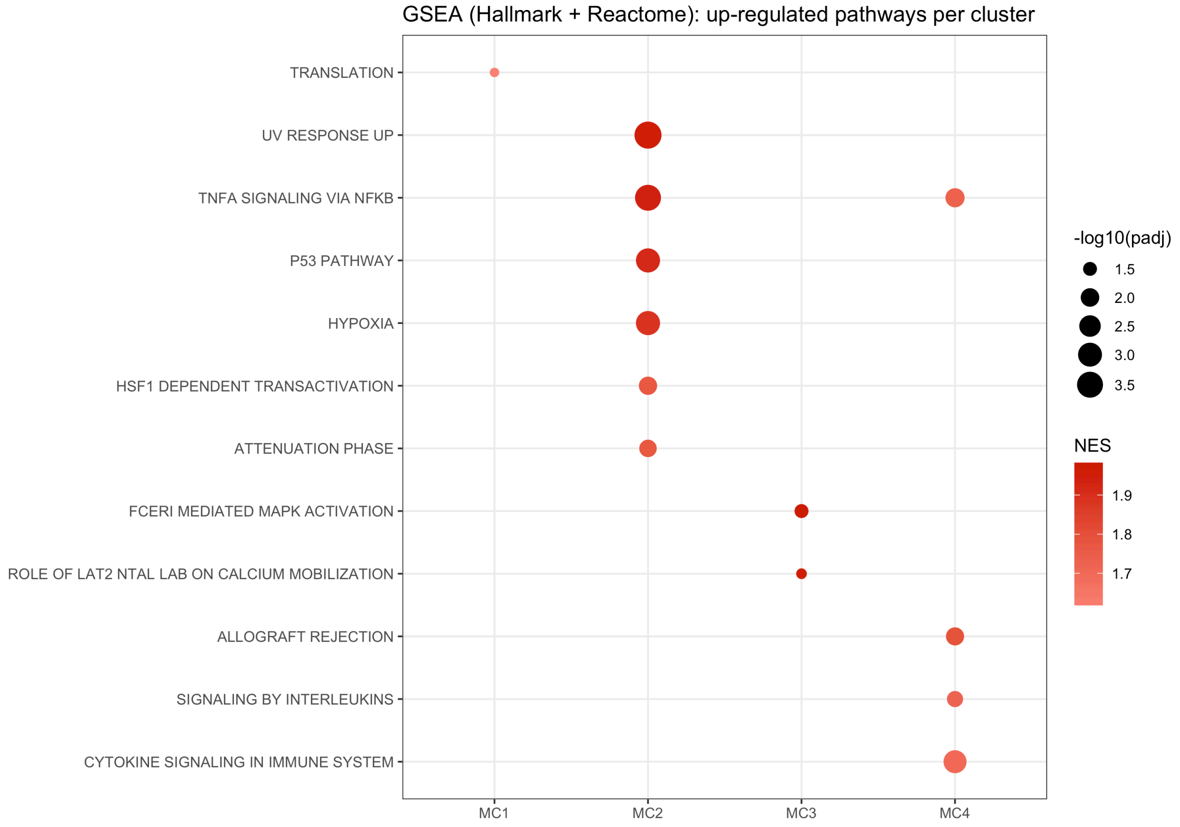


**Figure E3.** Gene set enrichment analysis (GSEA) per mast-cell cluster (fgsea; MSigDB Hallmark + Reactome). Dot colour, normalized enrichment score (NES); size, −log10 adjusted p-value.

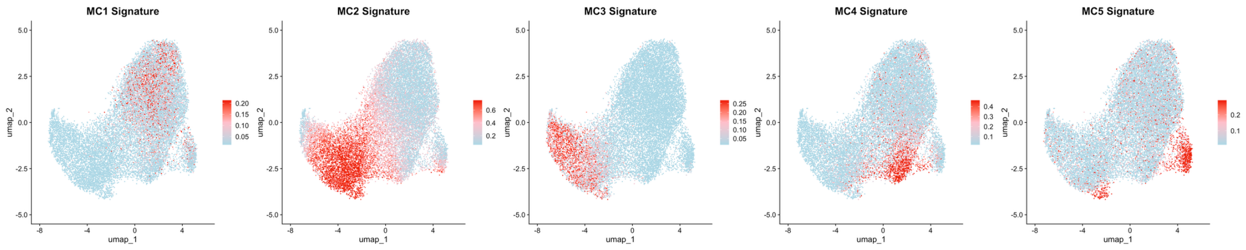


**Figure E4.** MC1–MC5 cluster signatures scored on our own UMAP (positive control).

### Online Repository Tables

**Table E1.** Per-donor quality-control thresholds.

| **Donor** | **Cell Ranger** | **Mito % (max)** | **nCount_RNA (min–max)** | **nFeature_RNA (min–max)** |
|---|---|---|---|---|
| 1 | v5.0.1 | 13 | 3,000–17,000 | 2,000–4,000 |
| 2 | v7.0.1 | 14 | 1,500–9,400 | 600–2,800 |
| 3 | v7.0.1 | 18 | 1,800–10,000 | 800–3,000 |
| 4 | v7.1.0 | 8 | 2,600–9,000 | 900–2,500 |
| 5 | v7.1.0 | 9 | 2,000–6,000 | 950–2,100 |

**Table E2.** Marker genes for clusters MC1–MC5. Genes were identified with Seurat FindAllMarkers, comparing each cluster with all remaining cells combined (MAST hurdle model, with donor as a latent variable). For each cluster the 50 genes with the highest average log2 fold change are listed, in order of p value (10 genes for MC5). Because each cluster is tested against all other cells pooled rather than against individual clusters, a gene can be significant here while not differing significantly between two particular clusters. The last two columns mark genes shown in Figure 2B and genes forming the cluster signatures used in Figure 6B.
